# Generation of a SARS-CoV-2 Replicon as a Model System to Dissect Virus Replication and Antiviral Inhibition

**DOI:** 10.1101/2020.12.12.422532

**Authors:** Xi He, Shuo Quan, Min Xu, Silveria Rodriguez, Shih Lin Goh, Jiajie Wei, Arthur Fridman, Kenneth A. Koeplinger, Steve S. Carroll, Jay A. Grobler, Amy S. Espeseth, David B. Olsen, Daria J. Hazuda, Dai Wang

## Abstract

SARS-CoV-2 research and antiviral discovery are hampered by the lack of a cell-based virus replication system that can be readily adopted without biosafety level 3 (BSL-3) restrictions. Here, the construction of a non-infectious SARS-CoV-2 reporter replicon and its application in deciphering viral replication mechanisms and evaluating SARS-CoV-2 inhibitors are presented. The replicon genome is replication competent but does not produce progeny virions. Its replication can be inhibited by RdRp mutations or by known SARS-CoV-2 antiviral compounds. Using this system, a high-throughput antiviral assay has also been developed. Significant differences in potencies of several SARS-CoV-2 inhibitors in different cell lines were observed, which highlights the challenges of discovering antivirals capable of inhibiting viral replication *in vivo* and the importance of testing compounds in multiple cell culture models. The generation of a SARS-CoV-2 replicon provides a powerful platform to expand the global research effort to combat COVID-19.

## INTRODUCTION

The current pandemic of coronavirus disease 2019 (COVID-19) caused by the newly emerged coronavirus, severe acute respiratory syndrome coronavirus 2 (SARS-CoV-2) has led to more than 62 million infections and 1.46 million deaths as of 30 November 2020 (*1*). In response to this public health emergency, an unprecedented swift and concerted effort has been launched globally to identify safe and effective therapeutics and vaccines against the rapidly spreading virus. Presently, about 2000 SARS-CoV-2 investigational programs and clinical trials have been registered (*2*). However, most of the treatment trials to date have focused on repurposing a limited number of existing antivirals approved for other indications, such as lopinavir/ritonavir, approved for treatment of HIV infection, or remdesivir, originally developed for treatment of Ebolavirus infection. Although compounds directly targeting SARS-CoV-2 are urgently needed, discovery and development of such compounds typically requires a decade of dedicated effort. One of the major bottlenecks is that the replication of the SARS-CoV-2 is yet fully understood, and assays used to study viral genome replication and test SARS-CoV-2 compounds must be performed in BSL-3 laboratories which are not readily available to most researchers.

To address the need for a broadly accessible and robust cell-based SARS-CoV-2 replication platform, a SARS-CoV-2 replicon was constructed. In this replicon, the spike (S) protein open reading frame (ORF) was deleted and replaced with a gene encoding a firefly luciferase (Luc) and green fluorescence (GFP) fusion protein. The sequences of envelope (E), membrane (M) and their intergenic region were also deleted and replaced with a neomycin-resistance gene (Neo). The replicon maintained all the genes and genetic elements necessary for replicating the full-length and subgenomic (sg) RNAs. As a result, the system can be used to study many aspects of virus life cycle, such as genome replication, gene expression and virus-host interactions. Deletion of the structural genes rendered the replicon defective in producing progeny virions, thereby enabling safe handling under BSL-2 conditions.

Another advantage of the transient replicon system over traditional live virus-based models is that the replicon RNA can be introduced to a variety of cell lines, allowing for assessment of viral replication under more physiologically relevant conditions. SARS-CoV-2 preferentially targets the respiratory tract, but may also spread to other organs and tissues such as heart, liver, brain, and kidneys (*3*). In infected tissues, the virus has been detected in multiple cell types (*3, 4*). However, in tissue culture very few cell lines are permissive to productive SARS-CoV-2 infection (*5*). As a result, most SARS-CoV-2 replication analysis and compound testing so far have been performed in Vero E6 cells which may not fully recapitulate the complexity of infections *in vivo*.

In addition, a high throughput antiviral assay has been developed using the described replicon system. Under those assay conditions, the sensitivities of viral replication to SARS-CoV-2 inhibitors were found to be cell line dependent, suggesting antiviral compounds may have different potencies against the virus infecting different cell types *in vivo*. Therefore, the replicon system also provides a valuable tool to screen, test and optimize antiviral compounds and to elucidate their mechanism of action.

## RESULTS

### Construction and characterization of SARS-CoV-2 replicon

A bacterial artificial chromosome (BAC) system was employed to construct the SARS-CoV-2 replicon. As the full-length SARS-CoV-2 replicon is ∼28k nt, its cDNA, together with the flanking regions, was synthesized as five fragments with overlapping ∼30 base pairs and subsequently assembled into a BAC vector. The replicon was also linked to a T7 minimal promoter upstream of the 5’ untranslated region (UTR), and a cassette containing a 26 nucleotide poly(A), a hepatitis D virus ribozyme (Rbz) and a T7 terminator at the 3’ end to yield precise viral genome terminus (**Fig.1A**).

To produce the replicon RNA, the BAC containing the replicon cDNA was linearized by restriction enzyme digestion and then transcribed *in vitro* by T7 RNA polymerase as described in the Materials and Methods. The 5’ end of the synthesized RNA was capped by an anti-reverse cap analog (ARCA). The replicon RNA was electroporated into indicated cell lines, and amplification of the replicon RNA was quantified by the expression of GFP or luciferase reporter genes. A similar *in vitro* transcription strategy has also been successfully used in the coronavirus reverse genetics systems to rescue recombinant SARS-CoV-2 viruses (*6-8*).

The replication of *in vitro* transcribed RNA was initially examined in 293T cells. According to the coronavirus replication model (*9*), the viral genome is copied by replication-transcription complex, both continuously to produce minus-strand templates for genome RNA synthesis and discontinuously to produce minus-strand templates for sg mRNA synthesis. These two types of RNAs have also been reported recently in the SARS-CoV-2 infected cells (*10*). The structural proteins such as S, E, M and N are encoded by sg mRNAs (**Fig. 1A**). Therefore, GFP in the replicon RNA transfected cells is also likely expressed by the sg mRNAs. To confirm the synthesis of the Luc/GFP sg mRNAs, RT-PCR analysis was performed on the transcripts in the electroporated cells. The forward primer was positioned in the leader sequence of 5’ UTR, and the reverse primer was situated in the Luc/GFP coding region (**Fig.1A**). This resulted in a primer pair only amplifying Luc/GFP sg mRNA. Consistent with the GFP expression, the Luc/GFP sg mRNA was detected as early as 12 h post electroporation (**Fig.1B**). The presence of the Luc/GFP sg mRNA also indicated that the *in vitro* transcribed replicon RNA produces a functional replicase complex.

**Fig. 1.**
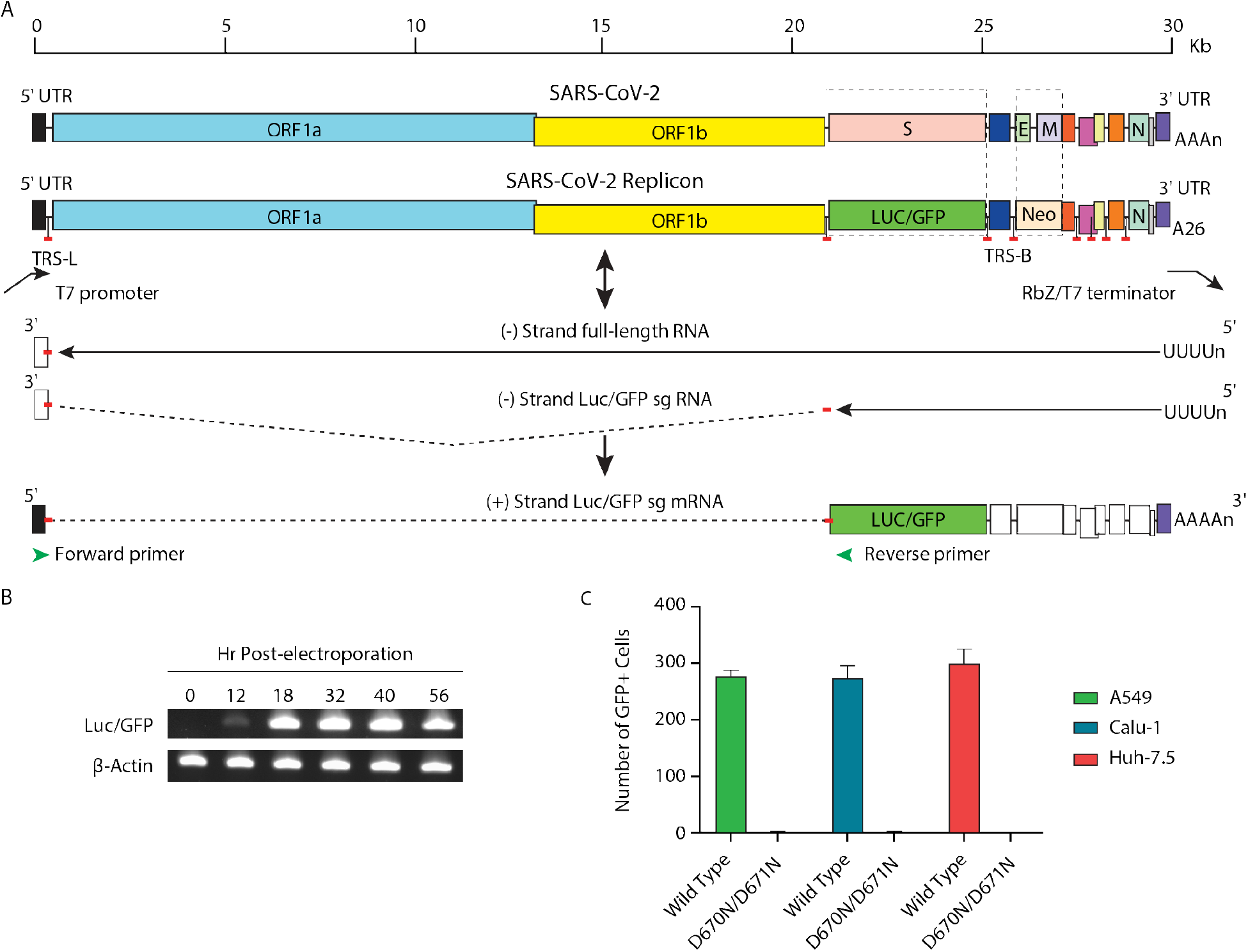
Construction and transcriptional analysis of SARS-CoV-2 replicon. **(A)** Schematic diagram of reporter replicon genome structure, replication and sg mRNA production. The S and E/M ORFs in the wild type viral genome were deleted and replaced with Luc/GFP and Neo genes, respectively. The full-length replicon cDNA was flanked by a T7 promoter and a polyA/HDV ribozyme and T7 terminator cassette. The *in vitro* transcribed replicon RNA is copied by replicase complex to produce genomic or sub-genomic sized negative stranded RNAs (solid lines). The negative stranded sg RNAs are used as templates to produce the sg mRNAs for expression of reporter and structural proteins. The sg mRNAs consist of the leader at 5’ UTR of the genome and mRNA body sequences joined by a short and conserved sequence motif, the transcription-regulating sequence (TRS). The untranscribed regions (dashed lines), TRS leader (TRS-L) and TRS body (TRS-B) are also indicated. **(B)** Detection of the sg mRNA expressing Luc/GFP reporter protein. Total RNA was collected and purified at indicated times from 293T cells electroporated with T7 polymerase transcribed replicon RNA. The mRNAs expressing Luc/GFP or β-actin were examined by RT-PCR. **(C)** GFP reporter signals produced by the wild type and RdRp mutant replicons. The A549, Calu-1 and Huh-7.5 cells were electroporated with T7 polymerase transcribed wild type replicon RNA or the RdRp D760N/D761N double mutant replicon RNA and transferred to 384 well plates. The number of GFP positive cells in each well were counted at 30 h post electroporation.

In addition to 293T cells, other cell lines such as A549, Calu-1 and Huh-7.5 cells were also demonstrated to support the replication of the SARS-CoV-2 replicon RNA, as evidenced by the GFP reporter gene expression in those cells following electroporation with the replicon RNA (**Fig. 1C**). To further investigate whether the reporter signals are generated by *de novo* synthesized RNAs, two point mutations, D760N and D761N, were introduced to RNA-dependent RNA polymerase (RdRp) in the replicon. The combination of the two mutations inactivates the RdRp and leads to a complete loss of native ribonucleotide incorporation (*11*). As expected, the mutant replicon RNA failed to express GFP in transfected A549, Calu-1 and Huh-7.5 cells (**Fig. 1C**). The data confirmed that reporter activities are a result of active replicon RNA replication and can be used as a readout to evaluate the effectiveness of potential inhibitors of enzymes required for replication of SARS-CoV-2.

### Inhibition of replicon replication by antiviral compounds

In order to demonstrate the sensitivity of the SARS-CoV-2 replicon to coronavirus inhibitors with different mechanisms of action (MOA), a series of pilot experiments were performed in 293T cells. The compounds chosen for these initial studies were GS-441524 and GC376. GS-441524 is the parent compound of remdesivir, a prodrug originally discovered as an antiviral against Ebola virus (*12*), specifically targeting the RdRp (*11*). Remdesivir has been approved by the US Food and Drug Administration (FDA) to treat hospitalized COVID-19 patients (*13*). Another notable compound GC376, a broad-spectrum inhibitor against 3C or 3C-like proteases that has previously been used to treat feline infectious peritonitis caused by feline coronavirus (*14*), is also reported to exert potent inhibition for the main protease (M^pro^ or 3CL^pro^) encoded by SARS-CoV-2 (*15*).

The 293T cells were electroporated with *in vitro* transcribed and capped full-length replicon RNA, and then incubated with increasing concentrations of GS-441524 or GC376 in a 384 well plate. Inhibition of replication was determined by reduction of the number of GFP positive cells or luciferase activities relative to DMSO control treated cells at 30 h post-electroporation. As shown in **Fig. 2A-D**, both compounds exhibited antiviral activities in a dose dependent manner, using either GFP or luciferase as a readout. The potencies of GS-441524 in inhibiting the SARS-CoV-2 replicon replication, determined by either % GFP signal or % luciferase activity, were 645 nM or 503 nM respectively. The potencies of GC376 in inhibiting the SARS-CoV-2 replicon replication, determined by either % GFP signal or % luciferase activity, were 22.0 nM or 29.2 nM respectively. The EC50 values measured by both reporters were comparable in these assays with the advantages of single cell-based GFP readouts for continuous monitoring of reporter gene expression, and of population-based luciferase readouts for fast assay development. The luciferase assay also confirmed that GFP expression reduction resulted from inhibition of replicon replication, not from quenching of GFP fluorescence. No cytotoxicity was detected for either compound in the range of concentrations tested (**Fig. 2E, F**).

**Fig. 2.**
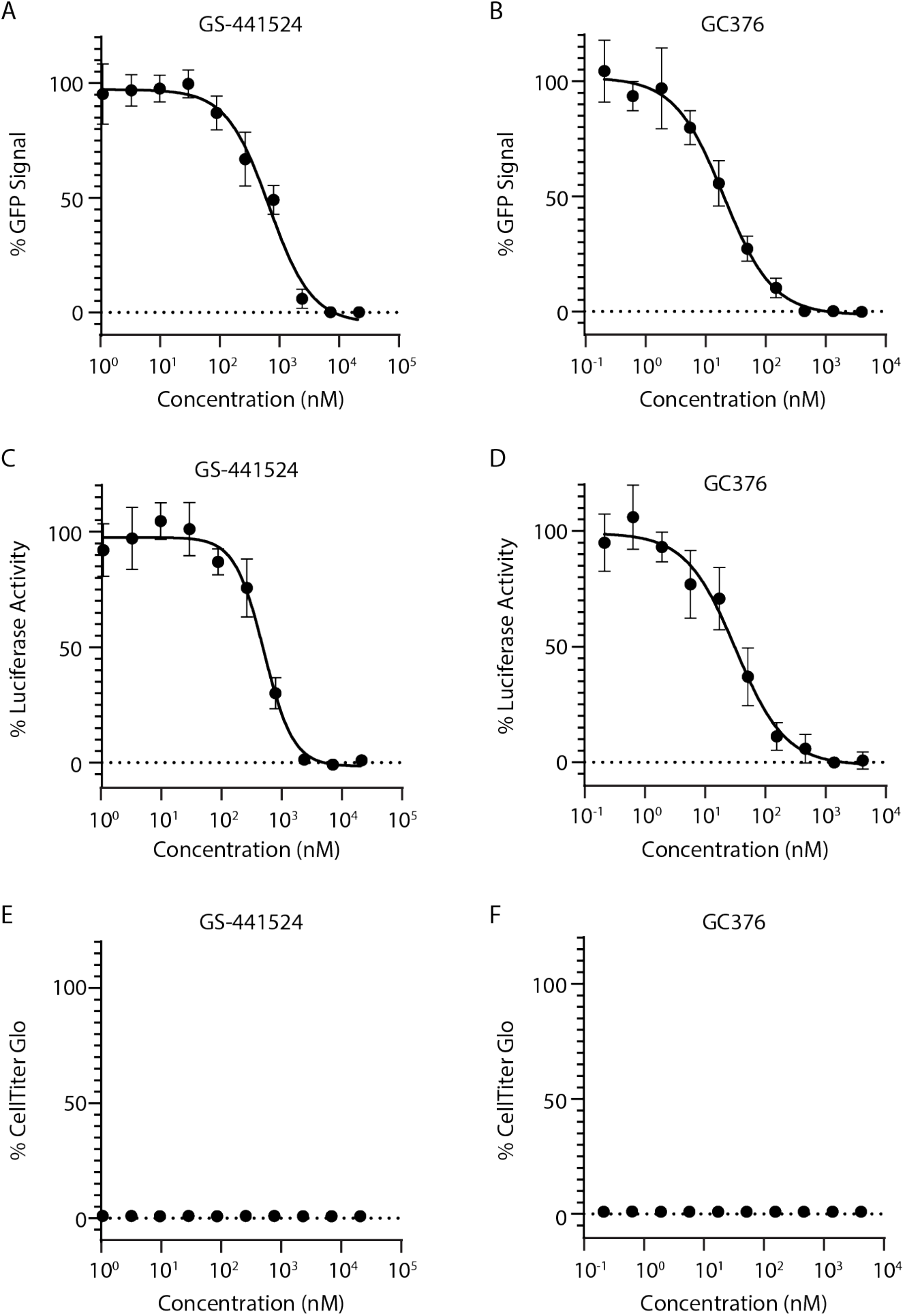
Dose dependent responses of SARS-CoV-2 replicon reporter activity to GS-441524 and GC376. The 293T cells were electroporated with T7 polymerase transcribed replicon RNA and incubated with GS-441524 or GC376 compounds at indicated concentrations. The activities were determined by % of GFP or luciferase signals comparing to DMSO treated cells **(A-D)**. The cytotoxicity of GS-441524 and GC376 to the cells were also examined **(E and F)**.

### Antiviral evaluation using SARS-CoV-2 replicon in different cell lines

Next we sought to determine the impact of cell lines on the potencies of a panel of SARS-CoV-2 compounds measured by the replicon assay. Those compounds have previously been shown to block SARS-CoV-2 replication at different stages of its life cycle.

As described above, all cell lines tested including 293T, Vero, Huh-7.5, Calu-1 and A549 could support replication of SARS-CoV-2 replicon. The compounds selected in this study were viral entry inhibitors, nafamostat and camostat (*16*) targeting transmembrane serine protease 2 (TMPRSS2), endosome trafficking inhibitors, relacatib (*17*) and apilimod (*17*) targeting cathepsin L and PIKfyve, respectively, polymerase inhibitors, remdesivir (*11*) and its parent compound GS-441524, the protease inhibitors GC373(*14, 18*), GC376 (*15*), PF-00835231 (https://doi.org/10.1101/2020.09.12.293498) and Boceprevir (*19, 20*).

Since the SARS-CoV-2 replicon RNA was introduced to the cells by electroporation, bypassing the natural entry pathways of the virus, as expected, the SARS-CoV-2 entry inhibitors were inactive in the replicon-based assay (**Table 1**). Also, due to the deletion of the structural proteins, S, E and M, this replicon assay is unlikely to be capable of identifying compounds that inhibit SARS-CoV-2 assembly or release.

**Table 1.**
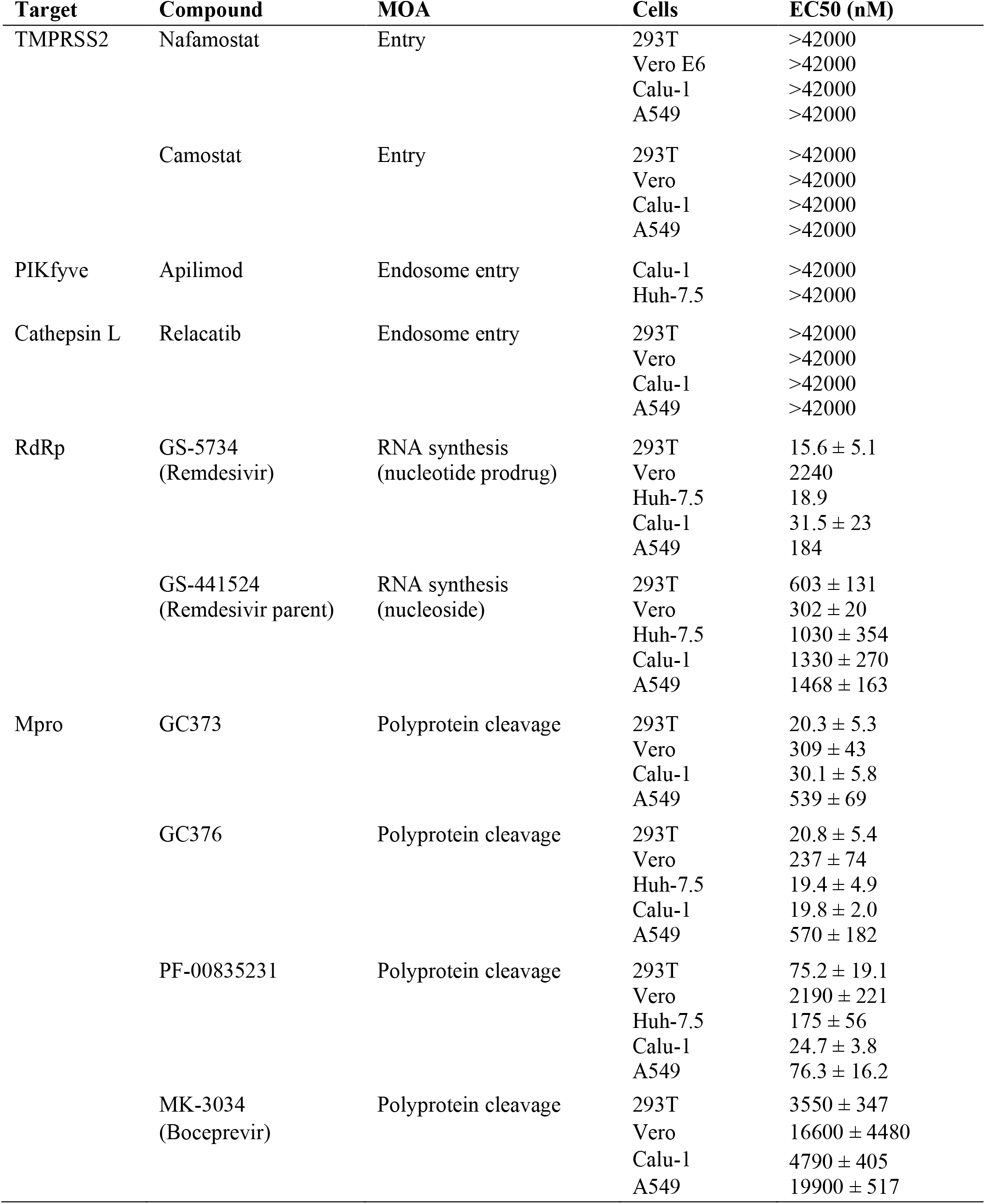
Antiviral activity against SARS-CoV-2 replicon.

Remdesivir and GS-441524 demonstrated antiviral effects in all cell lines, albeit to various degrees. The highest potency of remdesivir was found in 293T cells, with an EC50 value of 15.6 nM. In contrast, a much higher concentration of the compound is needed to achieve 50% inhibition in Vero cells (EC50 of 2240 nM), which could be attributed to inefficient cellular uptake and/or conversion of the nucleoside to the active triphosphate form. The reported EC50 values of remdesivir against wild type (wt) SARS-CoV-2 infection were 442 nM in ACE2 stably transfected A549 cells (https://doi.org/10.1101/2020.08.28.272880), and 770 to 1650 nM in Vero E6 cells (*21, 22*), consistent with the potencies of remdesivir measured in this study (Table 1). GS-441524 was relatively weaker in inhibiting the replicon activity in most cell lines compared to remdesivir.

The M^pro^ inhibitors, GC376, PF-00835231 and Boceprevir, also exhibited notable antiviral activities in different cell lines using the replicon-based assay. The sensitivity of the replicon to these compounds was also cell line dependent. GC376 and its parent compound GC373 (*14, 18*) were highly potent in 293T and Calu-1 cells, but only modestly active in A549 and Vero cells. In contrast, the EC50 value of PF-00835231 ranged from 32 to 175 nM in Calu-1, A549, and Huh-7.5 cells, and was 2190 nM in Vero cells. Boceprevir was the least potent among the M^pro^ compounds tested in this study.

Previously, it has been reported that Vero cells express high levels of the efflux transporter P-glycoprotein (p-gp) encoded by MDR1 or ABCB1 (*23*), which efficiently exports PF-00835231 and reduces the intracellular concentration of the compound (https://doi.org/10.1101/2020.09.12.293498). To determine the effect of p-gp on the activities of other M^pro^ inhibitors in Vero cells and the role of p-gp in other cell culture systems, the antiviral assay was then performed in the presence of p-gp inhibitors CP-100356 and elacridar, respectively. No antiviral activities were observed for either CP-100356 or elacridar at concentrations up to 2 µM when tested alone. Consistent with previous report (https://doi.org/10.1101/2020.09.12.293498), p-gp inhibitors dramatically increased the potency of PF-00835231 in Vero cells. There was also a slight increase of the activities of GC373, GC376 and Boceprevir in these cells. In contrast, the p-gp inhibitors did not cause any significant shift in the EC50 values of those compounds in A549 or Calu-1 cells.

### Adaptation of SARS-CoV-2 replicon-based assay for high throughput compound screening

The current replicon assay is performed in 384-well plates using automated liquid handling systems (**Fig. 3A**), therefore, the format is capable of screening large compound libraries. Results are available as early as 24 h following addition of drug. When conducting the high throughput screening of compounds, multiple factors may impact the assay performance or variability. Those contributing factors include bacmid integrity, RNA quality and overall cell health at the time of experiment. To further improve the assay consistency, a system using cryopreserved cell aliquots to screen and test compounds was developed. The EC50 values of the compounds determined using frozen cell aliquots were within error of the assay with those determined using freshly electroporated cells (**Fig. 3B**).

**Fig. 3.**
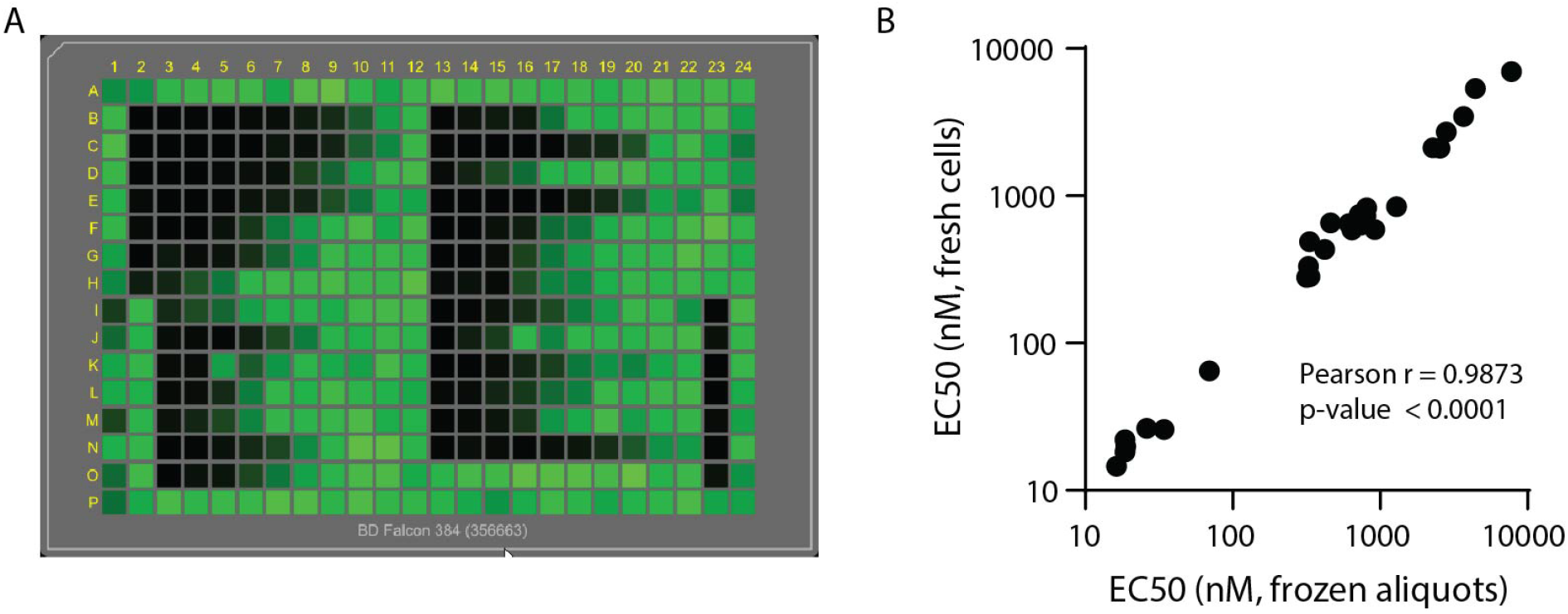
Adaptation of the SARS-CoV-2 replicon based assay to high throughput platform. **(A)** The plate view of GFP expression in replicon electroporated cells treated with compounds. The 293T cells were electroporated with T7 polymerase transcribed replicon RNA and incubated with control compounds in 384-well plates for 30 h. The plates were scanned using an Acumen eX3. **(B)** Correlation between the EC50 values measured using freshly or frozen electroporated cells. A total of 27 control compounds were tested using either fresh or cryopreserved 293T cells electroporated by SARS-CoV-2 replicon RNA as described in the Materials and Methods. The strength of correlation is determined by the Pearson test.

## DISCUSSION

Over the past decades, due in large part to the advances in reverse genetics, replicon systems have been established for a number of RNA viruses, such as hepatitis C virus, SARS-CoV-1, middle east respiratory syndrome coronavirus (MERS-CoV), dengue virus, West Nile virus, Zika virus, poliovirus enterovirus 71, hepatitis E virus, chikungunya virus and many others (*24*). For highly virulent viruses, the replicon assay system provides a safe and high throughput method to screen and evaluate compounds. The replicon-based assay also allows the identification of molecules that target proteins other than RdRp and M^pro^ whose functions are unknown or for which *in vitro* enzymatic assays are not yet understood, and elucidation of the compound mechanism of action via resistant selection (*25, 26*). Moreover, replicons make it possible to identify inhibitors against viruses that proved difficult to propagate in tissue culture, such as HCV (*27, 28*) and norovirus. Replicons have and will continue to play a crucial role in the discovery of new drugs against viral pathogens.

To construct and deliver the SARS-CoV-2 replicon, multiple strategies were explored in this study. Besides the current replicon design, a cytomegalovirus (CMV) immediate early gene promoter was evaluated to initiate the transcription of SARS-CoV-2 replicon, an approach reported previously for generation of SARS-CoV-1 replicon (*29*). The replicon cDNA under the control of a CMV promoter can be directly transfected into cells without the need of *in vitro* transcription. However, the replicon cDNA itself produced background GFP and luciferase signals, and the reporter activities were more variable from experiment to experiment.

We also attempted to establish a stable cell line that could maintain the replicon RNAs but to date have been unsuccessful. SARS-CoV-2 replicon might be toxic to the cells or cleared by intracellular innate immunity. Efforts to suppress or modify innate immunity to permit the tolerance of SARS-CoV-2 replicon are currently on-going.

The utility of the replicon system for studying SARS-CoV-2 genome replication and gene expression was demonstrated by the observations that the replicon is fully capable of producing sg mRNAs as seen in virus infected cells (**Fig. 1**). The expression of reporter genes could be inhibited by point mutations in RdRp or antiviral treatment. There were dramatic differences in potencies of the SARS-CoV-2 protease inhibitors in different cell lines. Multiple factors may contribute to the cell line dependent compound sensitivity. It is conceivable that the replicon may have different replication kinetics in different cells. High level expression of the protease may reduce the relative effectiveness of the compounds. The shift of potency is unlikely caused by the differences in electroporation efficiencies, since the A549, Calu-1 and Huh-7.5 showed similar number of cells expressing GFP in the assay (**Fig. 1C**). Another possible explanation is that rates of compound uptake, efflux and metabolism may be different in those cells, as shown in Vero cells (**Table 2**). Understanding the impact of cell lines on SARS-CoV-2 response to those compounds may also have an important implication in the clinical development of the drugs.

**Table 2.** Effect of p-gp inhibitors on the potency of compounds against SARS-CoV-2 replicon.

| Compound | EC50 (nM) |  |  |
| --- | --- | --- | --- |
|  | Vero | A549 | Calu-1 |
| GC373 | 309 ± 43 | 539 ± 69 | 30.1 ± 5.8 |
| GC373 + 0.5 µM CP-100356 | 99.2 ± 9.7 | 581 ± 16 | 29.4 ± 4.7 |
| GC373 + 2 µM CP-100356 | 64.2 ± 9.9 | 403 ± 30 | 20.9 ± 3.9 |
| GC373 + 0.5 µM Elacridar | 54.2 ± 0.4 | 656 ± 98 | 33.7 ± 8.4 |
| GC373 + 2 µM Elacridar | 35.8 ± 4.1 | 510 ± 147 | 27.0 ± 5.7 |
| GC376 | 237 ± 74 | 570 ± 182 | 19.8 ± 2.0 |
| GC376 + 0.5 µM CP-100356 | 63.8 ± 2.9 | 551 ± 3 | 9.16 ± 2.70 |
| GC376 + 2 µM CP-100356 | 44.0 ± 2.9 | 481 ± 83 | 9.42 ± 1.05 |
| GC376 + 0.5 µM Elacridar | 93.3 ± 15.7 | 631 ± 94 | 10.3 ± 2.04 |
| GC376 + 2 µM Elacridar | 46.4 ± 10.7 | 651 ± 124 | 9.40 ± 1.94 |
| PF-00835231 | 2190 ± 221 | 76.3 ± 16.2 | 24.7 ± 3.8 |
| PF-00835231+ 0.5 µM CP-100356 | 68.4 ± 5.7 | 61.0 ± 4.0 | 9.67 ± 1.10 |
| PF-00835231+ 2 µM CP-100356 | 17.3 ± 0.6 | 33.6 ± 2.3 | 9.59 ± 1.73 |
| PF-00835231+ 0.5 µM Elacridar | 25.0 ± 6.1 | 61.5 ± 17.2 | 12.4 ± 1.4 |
| PF-00835231+ 2 µM Elacridar | 16.3 ± 2.0 | 50.3 ± 9.3 | 9.68 ± 2.0 |
| Boceprevir | 16600 ± 4480 | 19900 ± 517 | 4790 ± 405 |
| Boceprevir + 0.5 µM CP-100356 | 7800 ± 1930 | 19800 ± 1480 | 4340 ± 1246 |
| Boceprevir + 2 µM CP-100356 | 4300 ± 1170 | 17900 ± 892 | 4050 ± 857 |
| Boceprevir + 0.5 µM Elacridar | 4000 ± 1290 | 16500 ± 699 | 4540 ± 570 |
| Boceprevir + 2 µM Elacridar | 4290 ± 1230 | 16800 ± 261 | 4080 ± 553 |
| CP-100356 | > 4670 | > 4670 | >5000 |
| Elacridar | > 4670 | > 4670 | > 8400 |

## MATERIALS AND METHODS

### Experimental design

The objective of this study is to develop a non-infectious, high throughput and cell line independent SARS-CoV-2 replicon based antiviral screening and testing system. The full-length cDNA of a SARS-CoV-2 reporter replicon was first synthesized and assembled into a bacterial artificial chromosome. Replicon RNA is produced in vitro using T7 polymerase and introduced to cells through electroporation. The amplification of the replicon RNA is then confirmed by the expression of sub genomic RNAs, and by the loss of the reporter activities as a result of RdRp inactivation. The replicon was further evaluated in an antiviral assay using known SARS-CoV-2 inhibitors. The impact of cell lines and efflux transporter on antiviral potencies of the compounds were also determined. The replicon-based assay has been automated and adapted to a 384 well plate format for high throughput compound library screening.

### Cell lines

Human embryonic kidney 293T (Clontech, 632180), human lung carcinoma A549 (ATCC, CCL-185) and human hepatoma Huh7.5 (Apath) cells were maintained in high-glucose Dulbecco’s modified Eagle’s medium (DMEM, Invitrogen, 10-013-CV); African green monkey kidney Vero (ATCC, CCL-81) and human lung epidermoid carcinoma Calu-1 (ATCC, HTB-54) cells were cultured in Eagle’s Minimum Essential Medium (ATCC, 30-2003) and McCoy’s 5a medium Modified (ATCC, 30-2007), respectively. All the media were supplemented with 10 % fetal bovine serum (FBS, Invitrogen, 10082147) and 1% penicillin/streptomycin. Cells were grown at 37°C with 5% CO2.

### Replicon construction

The replicon sequence is derived from the SARS-CoV-2 isolate Wuhan-Hu-1 (GenBank: NC045512). The replicon cDNA is placed under a T7 (5′ TAATACGACTCACTATA<u>G</u> 3′) promoter. The T7 RNA polymerase initiates transcription at the underlined G in the promoter sequence. Five fragments spanning the T7 promoter, full-length SARS-CoV-2, polyA/RbZ/T7 terminator (**Fig. 1A**), named F1 to F5, with ∼30 bps overlap were synthesized by Genewiz (South Plainfield, NJ). pSMART BAC vector (Lucigen) was digested with Not I. F1 and F5 fragments were digested with Mlu I. Equimolar amounts of linearized pSMART BAC vector, F1, and F5 were ligated using Gibson Assembly kit (NEB) according to manufacturer’s instruction, resulting in pSMART BAC F(1,5). Equimolar amounts of pSMART BAC F(1,5) digested with Aat II and Asc I, and F2 and F4 digested with Mlu I, were ligated using Gibson Assembly kit, resulting in pSMART BAC F(1,2,4,5). Finally, pSMART BAC F(1,2,4,5) digested with Aat II and Asc I, and F3 digested with Swa I were ligated together using Gibson Assembly Kit, resulting in the full-length non-infectious SARS-Cov-2 replicon construct pSMART BAC-T7-scv2-replicon.

### RdRp mutant replicon construction

The RdRp mutations were made to the replicon by BAC recombineering (*30*). An ampicillin (Amp) cassette was introduced to the RdRp immediately downstream of D761 residue, followed by Gibson assembly approach to replace the Amp cassette with the D760N and D761N mutations. pSMART BAC-T7-scv2-replicon was transformed into SW102 *E. coli* host expressing the bacterial phage λ Red genes *gam, bet and exo* which are controlled by *ts* repressor *c*I857. The Red enzymes are able to mediate homologous recombination between DNA fragments with overlapping sequences as short as 30 bp. The fragment containing an Amp cassette flanked by Asc I restriction sites and adjacent homologous sequence was amplified by PCR with primers 5’-TTTGTGAATGAGTTTTACGCATATTTGCGTAAACATTTCTCAATGATGATACTCTCT*GGC GCGCC*GGAACCCCTATTTGTTTATT-3’ and 5’-CATCTTAATGAAGTCTGTGAATTGCAAAGAACACAAGCCCCAACAGCCTGTAAGACT*G GCGCGCC*TTACCAATGCTTAATCAGTG-3’ using plasmid pcDNA3.1(+) as a template. The PCR fragment was then digested with Dpn I at 37 °C for 1 h and purified using the QIAquick PCR purification kit (Qiagen). To prepare the competent SW102 containing the pSMART BAC-T7-scv2-replicon, the E. coli host was first cultured in LB broth with 12.5 µg/ml of chloramphenicol for 4 h to reach log phase at 32 °C, followed by heat shock treatment at 42 °C for 15 minutes to induce expression of λ Red genes. After 3 washes with ice-cold water, 100 ng of PCR fragment containing Amp cassette was electroporated into the competent cells. After 2 h of recovery at 32 °C, the electroporated cells were plated onto LB plates containing 100 µg/ml carbenicillin and 12.5 µg/ml chloramphenicol, and then cultured overnight at 32 °C. Correct clones were identified by colony PCR followed by restriction enzyme digestion. The pSMART BAC-T7-scv2-replicon-Amp DNA was prepared using NucleoBond BAC 100 kit (Takara Cat# 740579). A fragment containing the RdRp D760N D761N mutation sequence was amplified by PCR using primers 5’-CGTAAACATTTCTCAATGATGATACTCTCTAACAATGCTGTTGTGTGTTTCAATAG-3’ and 5’-CAAAGAACACAAGCCCCAACAGCCTGTAAGACTGTATGCGGTGTGTACATAGC-3’. pSMART BAC-T7-scv2-replicon-AMP DNA was digested with Asc I and then ligated to the fragment containing the RdRp D670N/D671N mutations using Gibson Assembly kit resulting in pSMARTBAC-T7-scv2m-replicon.

### RNA *in vitro* transcription and electroporation

The replicon vectors (pSMART-T7-scv2-replicon or pSMART-T7-scv2m-replicon) were first linearized by Swa I (NEB), and then purified by phenol/chloroform extraction and ethanol precipitation. The fragments were dissolved in nuclease-free water. The mMESSAGE mMACHINE T7 ultra transcription kit (Invitrogen), which includes a cap analogue Anti-Reverse Cap Analogue (ARCA), was used to generate the replicon RNAs in the correct orientation from the linearized vector according to manufacturer’s instruction. Briefly, 100 µl of T7 transcription reaction, containing 4 µg of linearized BAC and 15 µl of extra GTP, was incubated at 37°C for 2.5 h to increase the length of the transcripts. After incubation, 5 µl of TURBO DNase was added and the reaction was incubated at 37°C for 15 min to digest DNA. The resulting RNA was purified by Monarch RNA cleanup kit (NEB).

The cells were harvested using TrypLE Select (Thermofisher Scientific), washed three times with PBS and resuspended in MaxCyte electroporation buffer to 1×10^8^ cells/ml. For 1×10^6^ of cells, 1 µg of replicon RNA was added into resuspended cells. The mixture was immediately transferred to a MaxCyte processing assembly (PA) (OC-100, OC-400 or R-1000) and electroporation was carried out by MaxCyte STX with the pre-loaded programs for transfecting different types of cells. After resting in PA for 20 min, transfected cells were transferred into 30 ml of complete DMEM (DMEM supplemented with 10% FBS and 1% penicillin-streptomycin). Following cell density adjustment (4 × 10^5^ cells per mL), the RNA-electroporated cells were plated into 384-well compound assay plates.

### Cryopreservation of electroporated cells

The 293T cells were prepared and electroporated as described above. After resting for 20 min, the cells were pelleted, resuspended in freezing media (45% FBS, 45% complete DMEM, 10% DMSO) at 2 × 10^7^ cells per ml, and frozen in a Mr Frosty Freezing container (ThermoFisher Scientific) at −70°C.

### Detection of sg mRNA expressing Luc-GFP fusion protein

Total RNAs were extracted from 1.2×10^6^ cells using RNeasy mini kit (Qiagen) at 0, 12, 18, 32, 40 and 56 h post electroporation. cDNA was synthesized and amplified using a SuperScript IV One-Step qRT-PCR kit (Thermofisher Scientific) according to manufacturer’s instructions. The PCR products were analyzed on an E-Gel 1% agarose (Thermofisher Scientific). The sequences of Luc/GFP primer set used to amplify the sg mRNA were: 5’-AGGTTTATACCTTCCCAGGT-3’ and 5’-TTTGTATTCAGCCCATAGCG-3’, and the sequences of β-actin primer set were 5′-GAGCACAGAGCCTCGCCTTT −3′ and 5′-TGGGGTACTTCAGGGTGAGG −3′.

### Compound testing

Compound plates were prepared by dispensing compounds (0.2 µl/well) dissolved in DMSO into wells of 384 well poly D lysine-coated plates (Corning 356663) or cell culture-treated, flat-bottom plates (Corning 3571) using an ECHO acoustic dispenser. Each compound was tested in a 10-point serial 3-fold dilution (final concentrations 42016 nM −2.1 nM). Controls included DMSO only and GC-441524 (final concentration 10 µM). A total of 20,000 transfected cells were added (50 µL/well) using Agilent Bravo to compound plates and the cells were maintained at 37°C/5% CO2/90% relative humidity. Reporter activities in the compound treated cells were quantified at 30 h post-transfection, by counting the number of green cells in each well using an Acumen eX3 scanner or by measuring luciferase activity with Steady-Glo Luciferase assay system (Promega) using EnVison (Perkin Elmer).

### Cytotoxicity assay

Compound toxicities in replicon-transfected cells were determined via the CellTiter-Glo (CTG) luminescent assay (Promega), where the number of viable cells is determined based on the quantitation of ATP. After the compound plates were scanned in the Acumen eX3, they were equilibrated to room temperature for 30 min. The CTG assay was carried out according to manufacturer’s protocol. Briefly, 30 µL of the pre-mixed CTG reagents was added into each well using the Agilent Bravo. After gently mixing, the reaction was incubated at room temperature for 10 min, and luminescent signals were measured using the EnVision (Perkin Elmer).

### Quantification and data analysis

All numerical data are presented as the mean ± SD (standard deviations). EC50 was determined by non-linear 4-parameter curve fitting using ActivityBase. Data presented in Fig. 1 were re-graphed in Prism (GraphPad).

